# Rational design of chemically responsive cytokines for cancer immunotherapy

**DOI:** 10.1101/2025.09.18.677079

**Authors:** Lucia Bonati, Stephen Buckley, Leo Scheller, Alessandro Minafra, Sandrine Georgeon, Tom Enbar, Sailan Shui, Li Tang, Bruno E. Correia

## Abstract

Cytokines are key signal mediators of the immune system, playing essential roles in regulating central immunological processes. Despite their unique ability to modulate the immune system, the translation of cytokine-based therapies to the clinic has been significantly hindered by severe toxicities resulting from the pleiotropy and off-target effects of many cytokines. Here, we present a general strategy for the design of switchable cytokines that enables precise control over cytokine activity using the clinically approved drug, Venetoclax. We rationally designed self-inhibited <u>Sw</u>itchable <u>I</u>nter<u>L</u>eukins (SwILs) by embedding the chemically controlled interface (Bcl-2: BIM-BH3) into the cytokine’s structure, resulting in conditional cytokines activated by Venetoclax.

We showed the broad applicability of this strategy across various cytokines, including IL-2, IL-15, and IL-10. *In vivo* studies with IL-15 demonstrated that the SwILs achieved Venetoclax-dependent tumor control comparable to that of the native cytokine. Additionally, we showed that this strategy can be expanded to respond to other tumor-intrinsic stimuli, such as tumor-specific proteases. Overall, SwILs offer control of cytokine activity, enhancing the safety and clinical applicability of cytokine-based therapeutics.

## Introduction

Adoptive transfer of tumor-specific T cells has had considerable success in cancer treatment^1^. However, such therapies are less effective against solid tumors, likely due to the immunosuppressive effects of the tumor microenvironment (TME)^2^. This poses a significant challenge, necessitating strategies to support tumor-reactive T cells within the complex environment of the TME.

Among the various agents being explored, stimulatory cytokines that can enhance activation, proliferation, and cytotoxic function by antigen-specific T cells stand out as promising candidates. However, severe treatment-derived side effects remain a major barrier to the clinical use of cytokine therapies^3^. This is largely attributable to the pleiotropic nature of cytokines, which leads to undesired binding and activation of off-target cells. Furthermore, native cytokines typically have a short serum half-life and narrow therapeutic window, severely limiting their therapeutic efficacy.

To enhance control over cytokine-based therapies, several efforts have been focused on developing conditionally activated cytokines that can be temporally or spatially controlled. Most approaches in this field have relied on exploiting tumor markers of various types, such as overexpressed antigens^4,5^, low pH^6–10^, altered ATP levels^11^, and elevated protease concentrations^12–18^. While these strategies can achieve a degree of spatial targeting, they lack precise control over the kinetics of the cytokine’s activity. Moreover, such approaches often assume homogeneity across tumor types, overlooking that different tumors vary in their molecular profiles and stimulus concentrations. This variability may result in inconsistent cytokine activation, leading to insufficient activation within the TME or undesired activity in non-tumor tissues. Consequently, the concept of externally controlling cytokine activation presents a compelling strategy.

Recent advances in computational and data-driven approaches have enabled the fast and efficient design of proteins, providing novel strategies to address key limitations in cytokine biology, including pleiotropy, suboptimal pharmacokinetics, and systemic toxicity. Protein design has been successfully employed to generate cytokine mimetics with biased receptor signaling^19^, to inhibit cytokine receptor pathways, and to achieve selective binding to specific receptor domain pairs^20–22^. For example, the Rosetta protein modelling suite has been utilized to engineer cytokine mimetics that engage only defined receptor complexes, such as an IL-2 mimetic selectively targeting the IL-2Rβγ_c_ complex independently of IL-2Rα, or an IL-4 mimetic that signals exclusively through the type I receptor complex^20,21^. This paradigm has been further expanded through the design of a split, conditionally active IL-2 variant that is functional only in the presence of multiple antigens^4^.

Previously, we reported the use of a computationally designed, chemically disruptable heterodimer (CDH) based on the Bcl-2:BIM-BH3 complex, where the BIM-BH3 motif was transplanted to the Lead Design 3 (LD3) protein using computational tools, resulting in an efficient ON- and OFF-switch platform for protein and cell-based therapeutics^23,24^.

In this study, we present a generalizable strategy utilizing the Bcl-2:BIM-BH3 CDH to achieve precise control over cytokine activity by selectively masking the cytokine’s receptor-binding site with a chemically responsive domain (Figure 1a). The selected masking domain is responsive to the clinically approved small molecule, Venetoclax (Figure 1b). We first engineered IL-15 and demonstrated both efficient inhibition in the absence of Venetoclax and subsequent activation upon drug addition. To further optimize ligand-induced switchability, we developed a series of IL-15 mutants with varying affinities between Bcl-2 and the BIM-BH3 motif, showcasing the capacity to tune the responsiveness of the Switchable InterLeukin (SwIL) (Figure 1c). Subsequently, we extended this approach to additional cytokines, IL-2 and IL-10, illustrating the modularity and generalizability of the system (Figure 1d). Finally, we adapted this engineering strategy to respond to an alternative stimulus, matrix metalloproteinases (MMPs), demonstrating the platform’s versatility (Figure 1e).

**Figure 1.**
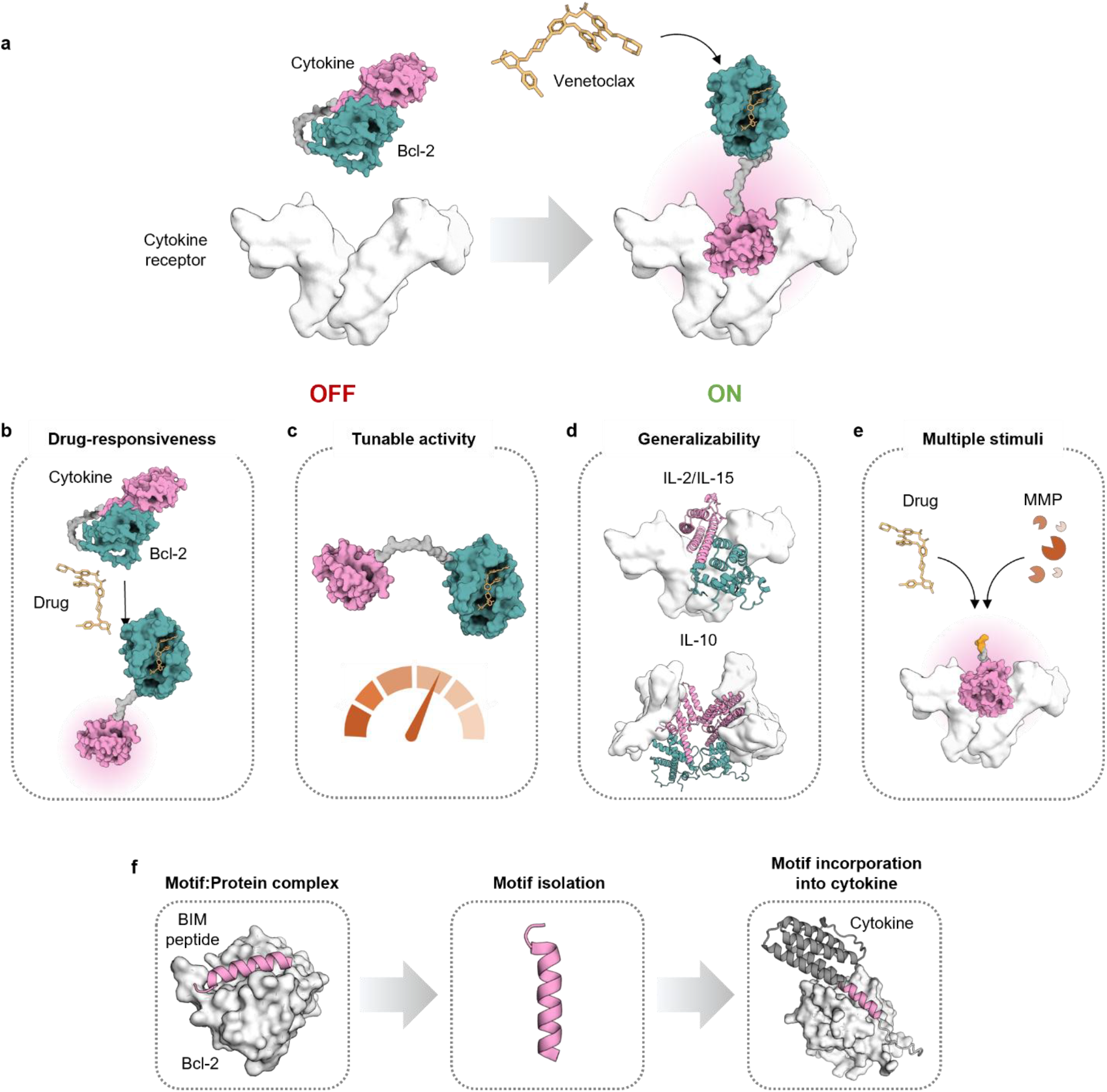
Overview of the switchable cytokine engineering strategy. **a**, The Bcl-2:BIM-BH3 CDH is incorporated into the cytokine’s structure to mask its receptor-binding site and inhibit its activity. Upon the addition of Venetoclax, the CDH is disrupted, activating the cytokine. This strategy provides control over the cytokine’s activity using Venetoclax (**b**) while allowing for CDH affinity tuning, which allows for differential masking efficiency and switching kinetics (**c**). This switchable cytokine platform can be generalized to cytokines of different families (**d**) and expanded to other stimuli, such as MMPs (**e**). **f**, After identification of a suitable motif:protein complex, we isolated the BIM peptide and incorporated it to the N- or C-termini of the cytokine of choice.

Altogether, we propose a novel strategy for the design of conditional cytokine activation triggered by either an exogenous small molecule or endogenous tumor stimuli, introducing a new dimension of control for cytokine-based therapeutics.

## Results

### Designer drug-controlled cytokines are functional *in vitro*

Conditional activity can be imparted to cytokines through the fusion of responsive domains, inhibiting cytokine-receptor interactions^12–14,17,25,26^. We hypothesized that fusing a CDH in close proximity to the cytokine receptor binding site would block cytokine-receptor interactions in the absence of the protein-protein interaction (PPI) disrupting drug. We reasoned that the recently developed CDH between human Bcl-2 (B-cell lymphoma 2; a transmembrane mitochondrial, nuclear, and endoplasmic reticulum membrane protein with anti-apoptotic activity) and the BH3 (Bcl-2 homology 3) domain of BIM (Bcl-2-interacting mediator of cell death; a pro-apoptotic molecule) would be an ideal system^27^. This CDH uses a short interaction motif as one half of the heterodimer, and is of particular interest because it is disrupted by the clinically approved inhibitor Venetoclax (ABT-199)^28^.

Leveraging the helical structure of most interleukins (ILs), we generated switchable constructs of Fc-fused human IL-15 (hIL-15/Fc). We incorporated the BIM-BH3 interaction motif and Bcl-2 to the termini of the cytokine, creating an intra-molecular CDH (Figure 1f). The spatial arrangement between Bcl-2 and the cytokine receptor was modulated by truncating or extending the termini of the protein, with elongation achieved through the insertion of alanine residues between the cytokine and the BIM-BH3 motif. We conducted an initial *in silico* screen by subjecting the rationally designed mutants to structural prediction via AlphaFold2 and selecting the variants predicted to exhibit steric hindrance with the receptor.

Next, we screened the selected variants, fused to an IgG1 Fc domain for extended half-life, in a HEK-293T cell-based assay (Supplementary Figure 1a). We transfected HEK-293T cells to express the selected mutants, and applied the resulting supernatants to reporter HEK-Blue^™^ IL-2R cells. These cells co-express the CD122 and CD132 (IL-2Rβ and IL-2Rγ) subunits of the IL-2 receptor and produce SEAP as a reporter for receptor engagement and JAK/STAT pathway activation. We screened a total of 10 switchable IL-15 (SwIL-15) designs, of which 5 induced SEAP expression in response to treatment with 1 μM Venetoclax (Figure 2a and Supplementary Figure 1b). Among these, mutant 5 (SwIL-15_mut5) demonstrated the highest fold-change between induced and non-induced conditions. This variant features a C-terminal BIM peptide (Figure 2d). SwIL-15_m5 was subsequently purified, and its expression and molecular weight confirmed by SDS-PAGE (Supplementary Figure 1e, h).

**Figure 2.**
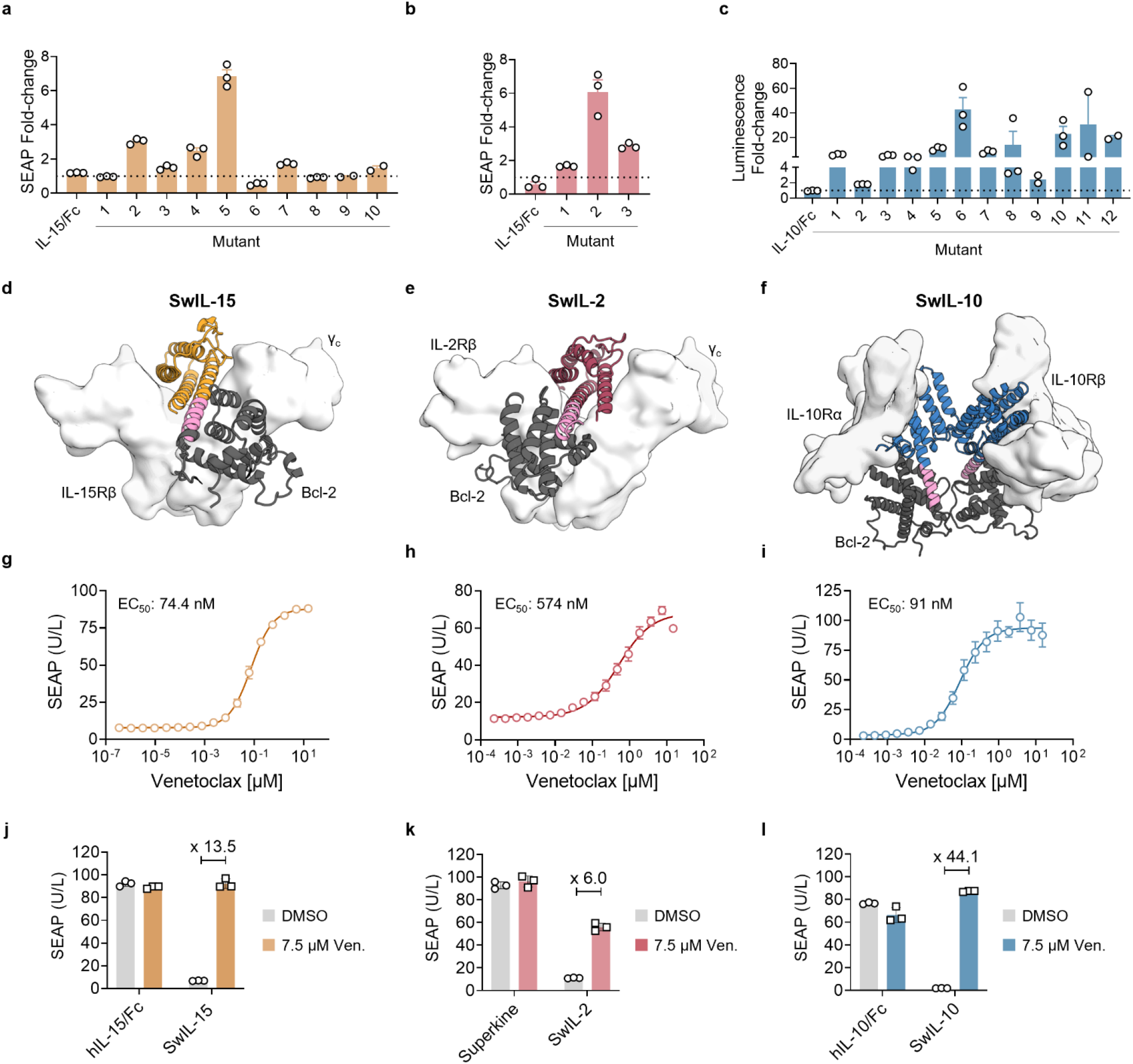
Screening of cytokine variants results in switchable mutants of IL-15, IL-2, and IL-10. **a-c**, Screening results for SwIL-15 (**a**), SwIL-2 (**b**), and SwIL-10 mutants (**c**). HEK-Blue^™^(**a, b**) or HEK293T cells expressing IL-10R and a pSTAT3-SEAP reporter (**c**) were treated with supernatants containing the switchable variants in the absence or presence of 1 μM Venetoclax. Shown are the fold-changes of reporter activity for induced/uninduced (1 μM Venetoclax/no Venetoclax) cells. **d-f**, Models of the selected best-performing cytokines in complex with their respective target receptors: SwIL-15_mut5:IL-15R (**d**), SwIL-2_mut2:IL-2R (**e**), and SwIL-10_mut10:IL-10R (**f**); Dark-grey: Bcl-2, pink: BIM peptide. **g-i**, Venetoclax-dependent dose-responses performed in HEK-Blue^™^induced with 5 nM SwIL-15 (**g**) and 1.5 nM SwIL-2 (**h**) or HEK293T cells expressing IL-10R and a pSTAT3-SEAP reporter induced with 5 nM SwIL-10 (**i**). SwIL-15_mut5 EC_50_ = 74.4 nM; SwIL-2_mut2 EC_50_ = 574 nM; SwIL-10_mut10 EC_50_ = 91 nM. **j-k**, SEAP expression in HEK293T cells expressing IL-2/15R treated with 5 nM of hIL-15/Fc or SwIL-15 (**j**), superkine or SwIL-2 (**k**) with or without 7.5 μM Venetoclax (Ven.). **l**, SEAP expression in HEK293T cells expressing IL-10R treated with 5 nM of hIL-10/Fc or SwIL-10 with or without 7.5 μM Venetoclax (Ven.). P-values of comparing induced and non-induced samples equal to p < 0.0001 in (**j**) and (**k**), and p = 0.0005 in (**l**). P-values comparing induced samples equal to p > 0.05 in (**j**), p < 0.001 in (**k**), and p < 0.01 in (**l**). Throughout, data represent the mean ± s.e.m., and p-values are determined by one-way Tukey’s test. Experiments were performed at least twice, with similar results. Representative data are shown.

We functionally characterized the design using HEK-Blue^™^ IL-2R cells exposed to the cytokine in the presence or absence of Venetoclax. The dose-response experiments revealed that increasing concentrations of Venetoclax progressively activated SwIL-15_m5, reaching activation levels in the same range of unmodified hIL-15/Fc and yielding an average EC_50_ of 74.4 nM (Figure 2g, j, Supplementary Figure 1b,f). In the absence of the drug, SwIL-15_m5 exhibited significantly low activity compared to unmodified hIL-15/Fc; however, upon treatment with Venetoclax, it achieved a ∼13.5-fold induction (Figure 2j).

To assess the generalizability of our engineering strategy, we applied a similar approach used for SwIL-15 to other cytokines, including human IL-10 and the IL-2 ‘superkine’, as they are key cytokines for cancer immunotherapy^29–32^. For both cytokines, we generated mutants by fusing the BIM peptide to either the N-or C-terminus. The IL-2 ‘superkine’ is an IL-2 mutein designed to exhibit increased affinity for the dimeric IL-2Rβγ_c_ receptor complex, thereby enhancing selective activation of effector T cells. Given its ability to elicit potent antitumor responses with reduced toxicity in *in vivo* mouse models, the IL-2 superkine was selected as a promising candidate for further engineering^32,33^. Moreover, the increased affinity of the IL-2 superkine for IL-2Rβγ_c_ facilitated screening in HEK-Blue^™^ IL-2R cells. Screening these variants identified inducible mutants with promising activity profiles for both switchable IL-2 (SwIL-2) superkine and IL-10 (SwIL-10) (Figure 2b, c and Supplementary Figure 1c, d).

Variants resulting in high fold-changes were purified and subjected to biochemical and functional characterization. Incorporation of the BIM peptide to the N-terminus of the IL-2 superkine produced the most effective inducible variant, resulting in an EC_50_ of 574 nM and a 6-fold increase in signal upon Venetoclax treatment (Figure 2e, h, k, Supplementary Figure 1f). Similar to SwIL-15, the SwIL-10 design showing the highest fold-changes between the induced and uninduced was engineered with a BIM peptide fused to the cytokine’s C-terminus, separated from the cytokine by a single alanine residue (Figure 2c, f, and Supplementary Figure 1d, g). This SwIL-10 variant exhibited no detectable activity in the absence of Venetoclax but showed a ∼44-fold induction upon drug treatment with an EC_50_ of 91 nM (Figure 2I, l, Supplementary Figure 1g). These findings suggest that incorporating the BIM peptide into more structurally complex interleukins, such as homodimeric cytokines, enables successful conditional activation. Collectively, the results demonstrate that embedding the BIM peptide within the helical structure of cytokines effectively suppresses their activity, which can be selectively restored in a Venetoclax-dependent manner.

### Tuning the affinity of the BIM Bcl-2 interaction modulates switching kinetics

While SwIL-15_mut5 showed small-molecule-dependent activation, we sought to engineer a cytokine with rapid activation kinetics to enable a fast switch-like behavior *in vivo*. However, after incubation of SwIL-15_mut5 with Venetoclax over varying time intervals, we assessed that SwIL-15_mut5 activation required overnight incubation with Venetoclax, indicating suboptimal, slow-switching kinetics (Supplementary Figure 2a). We hypothesized that the intramolecular interaction of the BIM peptide to Bcl-2 would further enhance the nanomolar affinity of the intermolecular interaction. Consequently, the increased affinity could potentially impede effective competition by Venetoclax, thereby limiting the disruption of the BIM:Bcl-2 interaction and reducing the cytokine activity. With these considerations, we aimed to engineer SwIL-15_m5 variants with enhanced switchability. To achieve that, we used Rosetta’s protein modeling framework to conduct *in silico* site-saturation mutagenesis (SSM) on all BIM peptide residues to highlight mutants with increased computed binding energy (ΔΔG) (Supplementary Figure 2b). This approach identified several point mutations (I120A, I127A, and F131A) with reduced binding affinity. Kinetics measurements of STAT5 phosphorylation of the identified mutants showed that the I120A mutation had markedly improved switching kinetics, achieving activation comparable to unmodified hIL-15/Fc only 20 minutes after the addition of Venetoclax (Figure 3a, b). In contrast, the I127A and F131A mutations showed only slight kinetic improvements and were, therefore, discarded (Supplementary Figure 2c-e). Armed with this toolbox of fast-switching SwIL-15 proteins, we next turned our attention to the application of these cytokine switches to control cytokine signaling in primary cells.

**Figure 3.**
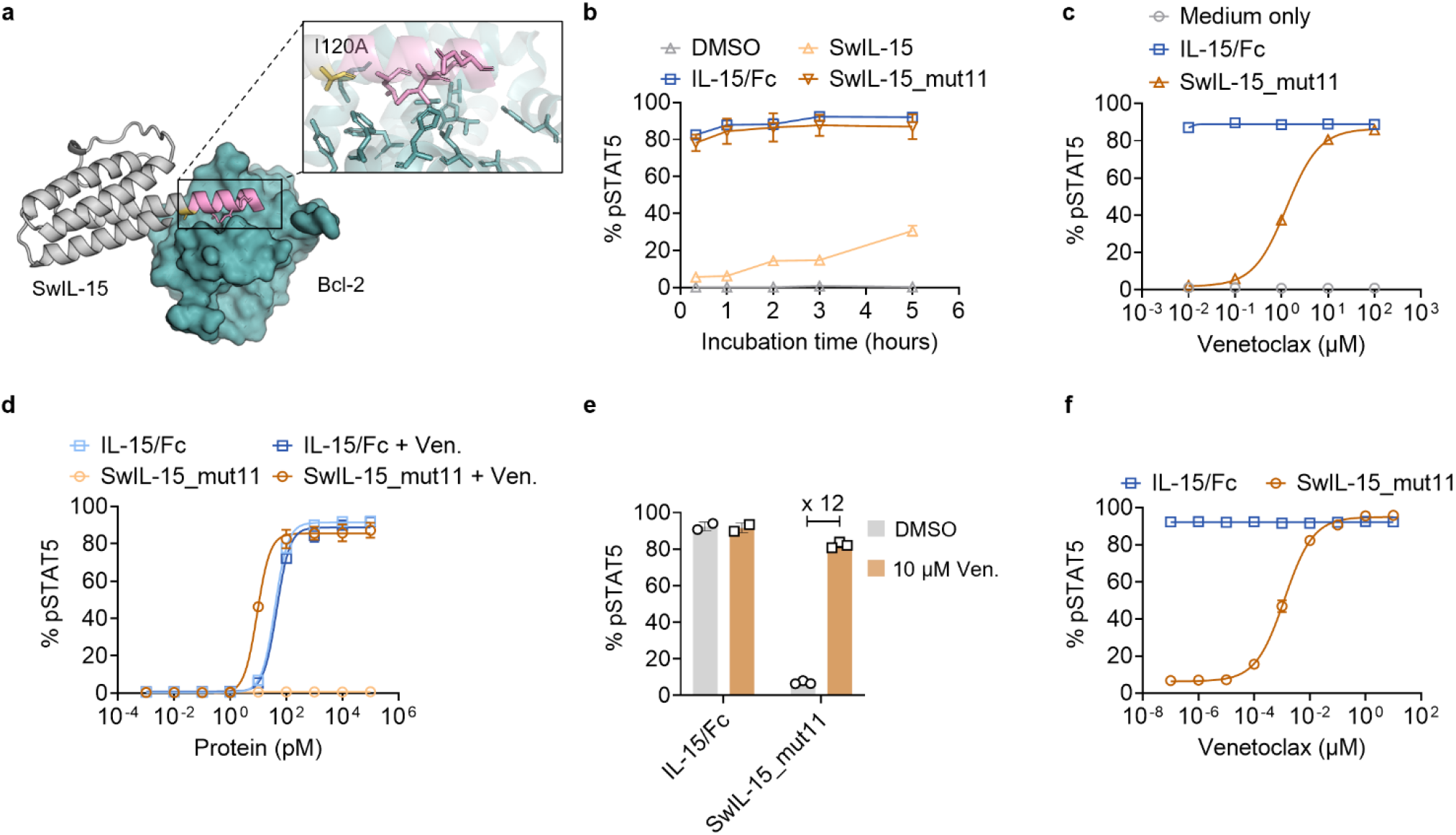
Switchable IL-15_mut11 fully regains activity upon treatment with Venetoclax. **a**, Model incorporating the beneficial mutations that decreased the affinity of the designed SwIL-15 to Bcl-2, thus improving switching kinetics. **b**, Switching kinetics analysis. 5 nM IL-15/Fc, SwIL-15_mut5, and SwIl-15_mut11 were first incubated with 10 μM Venetoclax for 0.33, 1, 2, 3, and 5 hours at 37°C and then used to treat pre-activated primary mouse T cells. **c**, Venetoclax-dependent dose-response of phosphorylated STAT5 (pY694) with Venetoclax-treated SwIL-15_mut11 or IL-15/Fc in pre-activated primary mouse T cells. EC_50_ = 457 nM. **d**, Protein-dependent dose-response of pSTAT5 with 10 μM Venetoclax-treated and untreated SwIL-15_mut11 and IL-15/Fc in pre-activated primary mouse T cells. SwIL-15 I120A EC_50_ = not detectable, SwIL-15_mut11 + Ven. EC_50_ = 9.2 pM, IL-15/Fc EC_50_ = 39.5 pM, IL-15/Fc + Ven. EC_50_ = 46.0 pM. **e-f**, pSTAT5 induction in pre-activated human primary T cells treated with 5 nM of Venetoclax-treated or untreated cytokines (**e**) and Venetoclax-dependent dose-response (5 nM of IL-15/Fc or SwIL-15_mut11) (**f**). Experiments were performed at least twice, with similar results. Representative data are shown. Throughout, data represent the mean ± s.e.m..

### SwIL15_mut11 activates primary T cells under the control of Venetoclax

Next, we sought to explore the functional activity of SwIL-15 I120A (from now on referred to as SwIL-15_mut11) in primary mouse T cells by measuring phosphorylation of STAT5. SwIL-15_mut11 demonstrated robust activation in a Venetoclax dose-dependent manner (Figure 3c). Additionally, SwIL-15_mut11 displayed a dose-response profile closely matching that of its unmodified counterpart (Figure 3d). Subsequently, we investigated the activity of the engineered cytokine in primary human T cells. In the absence of Venetoclax, SwIL-15_mut11, but not the unmodified version, showed strong signaling inhibition, while drug treatment restored activation levels comparable to those of unmodified hIL-15/Fc Figure 3e-f). Together, these findings demonstrate that SwIL-15_mut11 showed a switchable behavior when applied to both primary mouse and human T cells and maintained correct IL-15 functionality.

### Venetoclax-controlled SwIL-15_mut11 is effective in tumor models

As IL-15 has been shown to elicit an antitumor immune response *in vivo*, we decided to further characterize *in vivo* activity of SwIL-15_mut11^34^. First, we evaluated the pharmacokinetics in naïve mice by injecting equimolar amounts of IL-15/Fc and SwIL-15_mut11 intraperitoneally (i.p.) and collecting blood samples (Supplementary Figure 3a). We also tested different administration routes where SwIL-15_mut11 was injected i.p. and Venetoclax subcutaneously (s.c.) to simulate the absorption of epithelial cells. We observed no significant difference between the half-life of the unmodified IL-15/Fc and the masked SwIL-15_mut11, ranging from 6.6 to 7 hours. However, in mice injected with the SwIL-15_mut11 and Venetoclax, we observed a decrease in protein half-life of about 3 hours, resulting in a half-life of 3.7 hours (Supplementary Figure 3b-c). Given that experiments evaluating the effects of Venetoclax on tumor growth did not demonstrate a direct impact on tumor progression (Supplementary Figure 3d-e), we then assessed the antitumor efficacy of SwIL-15_mut11 in the presence or absence of Venetoclax in a subcutaneous B16F10 melanoma model. We treated B16F10-tumor bearing mice with adoptive cell transfer (ACT) of activated antigen-specific Pmel CD8^+^ T cells, combined with injections of IL-15/Fc or SwIL-15_mut11 (i.t., 200 pmol) with or without daily injections of Venetoclax (s.c., 25 mg/kg) (Figure 4a). SwIL-15_mut11 treatment alone failed to extend the survival of tumor-bearing mice compared to the control group (Figure 4b, d), demonstrating that the activity of SwIL-15_mut11 is blocked without the activating drug.

**Figure 4.**
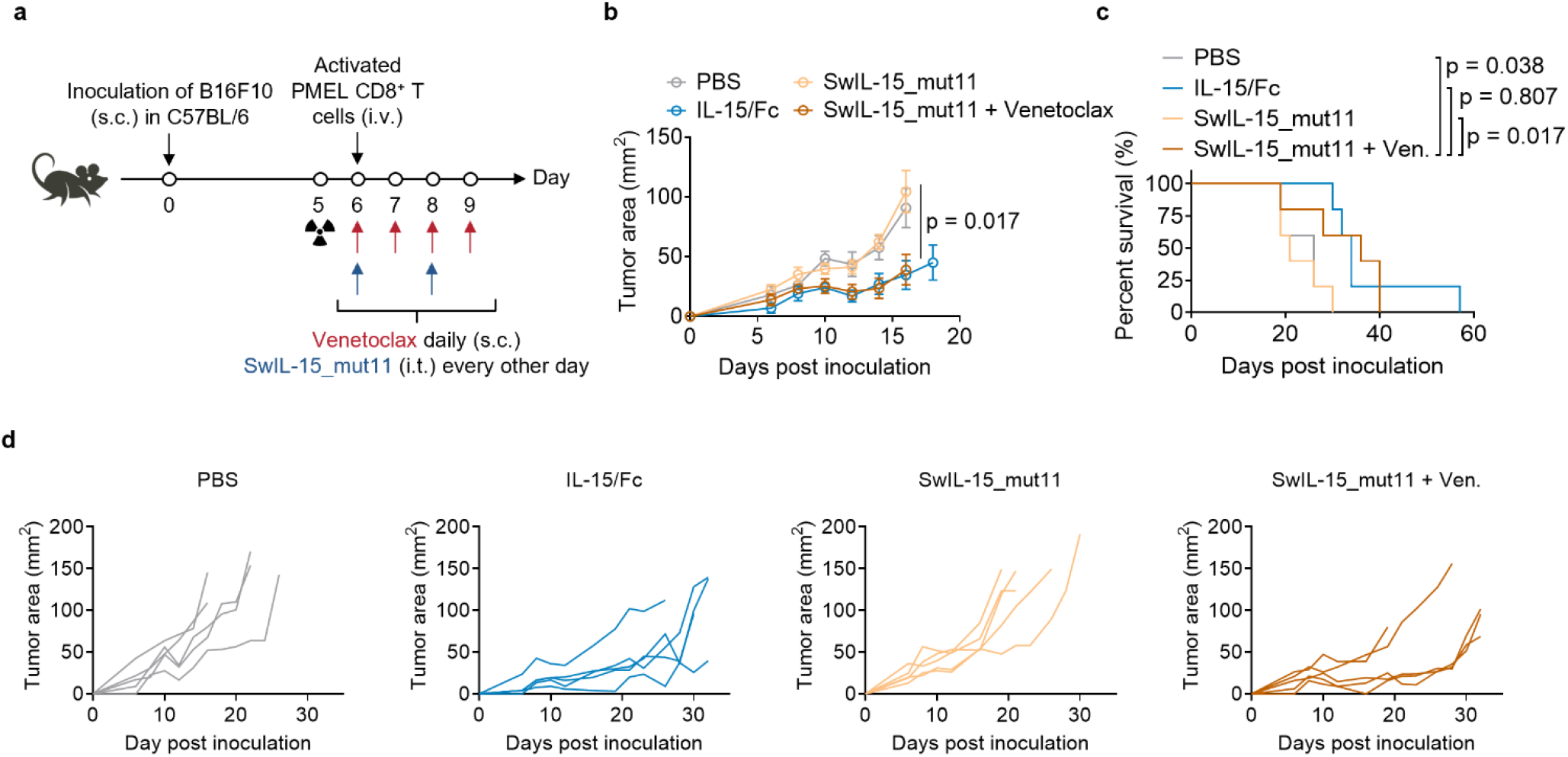
SwIL-15_mut11 induces a strong antitumor response equivalent to the native cytokine. **a-d**, C57BL/6 mice were inoculated subcutaneously (s.c.) with B16F10 melanoma tumor cells (3 × 10^5^), lymphodepleted by X-ray irradiation (4 Gy), and received adoptive transfer of CD8^+^T cells (5 × 10^6^) on day 6. Tumor-bearing mice were treated with IL-15/Fc (100 pmol/injection) and or SwIL-15_mut11 (100 pmol/injection) every other day with or without daily Venetoclax administration (25 mg/kg) as outlined in the experimental scheme (**a**). Mice receiving injections of PBS only serve as controls (n=5 animals per group). Shown are tumor growth curves (**b**), survival curves (**c**), and individual tumor growth curves. P-values were determined by one-way Tukey’s test (**b**) and log-rank test (**c**).

On the other hand, the combination treatment of SwIL-15_mut11 and Venetoclax significantly decreased tumor growth and extended mice survival in approximately 60% of treated animals, resulting in antitumor efficacy levels comparable to mice treated with ACT and IL-15/Fc (Figure 4b-d). Moreover, the IL-15/Fc intervention did not cause overt side effects, indicated by body weight drop (Supplementary Figure 3f-g). Overall, these results demonstrate that upon triggering by Venetoclax, the SwIL-15_mut11 is successfully activated *in vivo* and leads to a sustained tumor growth control comparable to its unmodified counterpart, serving as an *in vivo* proof of concept for switchable biologics.

### Harnessing tumor microenvironment cues to control SwILs activity

Drug-mediated control of cytokine activity offers a powerful approach to fine-tune activity levels and treatment duration. However, spatial targeting remains crucial, and the concept of engineering cytokines that are selectively active within the tumor microenvironment while sparing healthy tissues and circulation remains an unsolved problem. Therefore, we tested whether the same design principle could be adapted to respond specifically to tumor-specific stimuli. A promising stimulus is represented by MMPs, which are reported to be found in high concentrations in different types of solid tumors, such as melanoma, breast, and colon cancer^35–37^. We hypothesized that incorporating an MMP-cleavable linker between SwIL-15_mut11 and Bcl-2 would make the SwILs active in the presence of MMPs (Figure 5a). To design an MMP-responsive SwIL-15 (MMP-SwIL-15), we selected three MMP-cleavable peptides and identified suitable linkers to flank the cleavable peptide sequence. Linker options were classified as rigid (EAAAK or PAPAP) or flexible (GS)_n_, and designs included exclusively flexible, exclusively rigid, or mixed linkers (Figure 5b). Following sequence generation, we conducted an *in silico* screening through Alphafold2 to evaluate structural folding and ensure that (1) the interaction site between Bcl-2 and the BIM peptide was preserved and (2) the cleavable sequence was adequately exposed for efficient MMP-mediated cleavage. The most promising candidates were expressed, purified, and further characterized through functional assays. We assessed the proteolytic cleavage and bioactivity of MMP-SwIL-15 variants by treating them with MMP-2 and measuring STAT5 phosphorylation in activated T cells. All tested designs exhibited MMP inducible activation to differing degrees, ranging from 1.6x (L3, Supplementary Figure 4a) to 59x (L1-v1) increases in activation (Figure 5c-d). Figure 5 Next, MMP responsiveness was assessed for three designs (L1-v1, L1-v4, and L2-v2) (Figure 5e and Supplementary Figure 4b). All three designs showed an MMP dose-dependent response, indicating that different MMP concentrations can regulate their activity. The L1-v1 variant showed the highest fold change in activation and was the most sensitive to lower concentrations of MMP. It also maintained an inhibited state in the absence of MMP, resulting in minimal background activity. Collectively, these findings demonstrate that incorporating a cleavable linker between SwIL-15 and Bcl-2 is a viable strategy to achieve MMP-specific cytokine activation. Furthermore, this approach underscores the broader potential of integrating a Bcl-2-binding peptide to modulate cytokine responsiveness in a targeted and adaptable manner.

**Figure 5.**
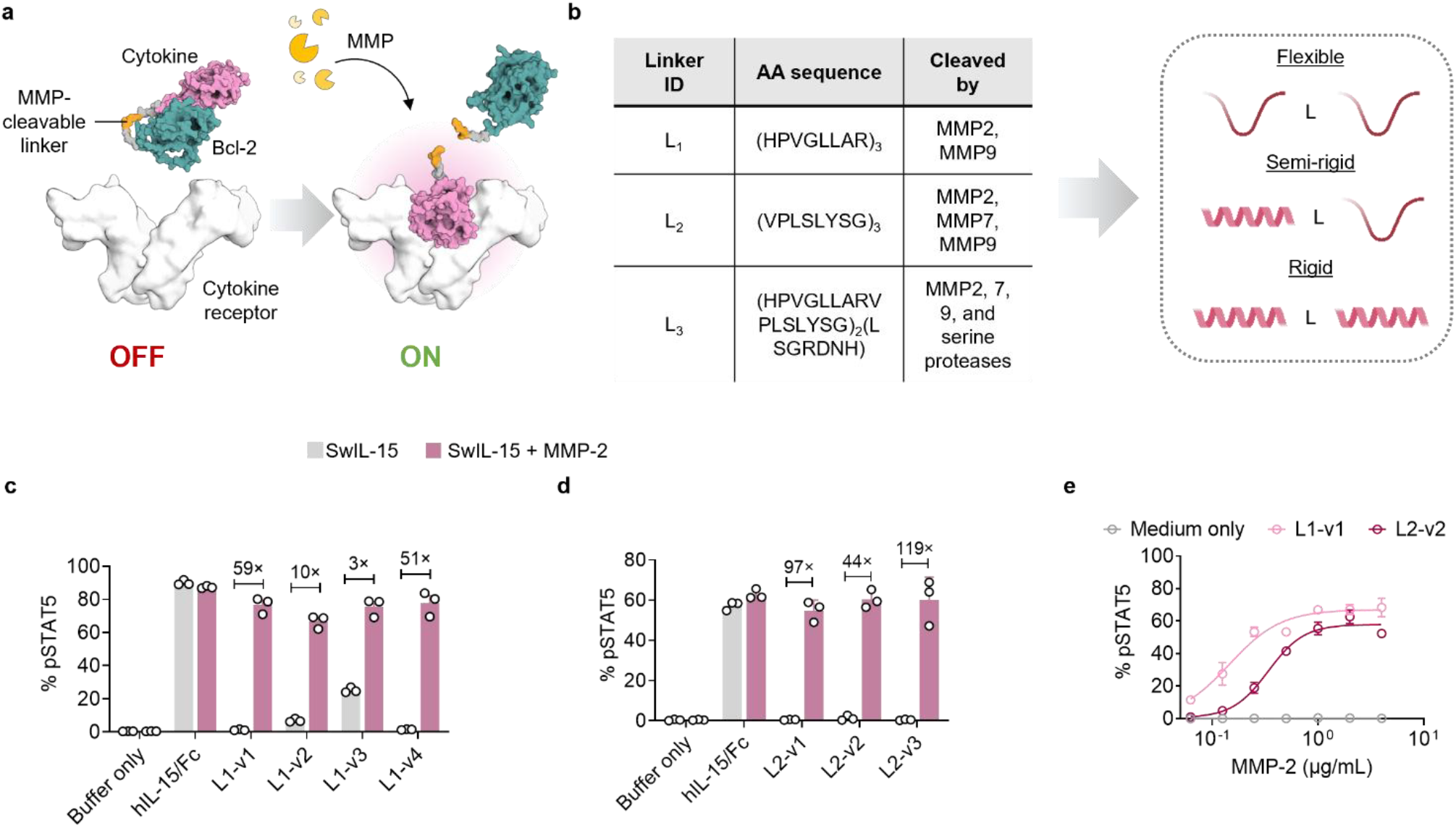
Switchable IL-15 system can be adapted to respond to tumor-intrinsic proteases. **a**, Schematic of masked SwIL-15 having incorporated an MMP-cleavable linker. In the absence of MMPs, Bcl-2 masks the receptor-binding site of IL-15 (no signaling), while in the presence of MMPs, Bcl-2 is cleaved, rendering the cytokine accessible to bind its cognate receptor. **b**, MMP-cleavable linker engineering strategy. (Left) Table of the amino acid (AA) sequences of linkers used to generate MMP-responsive SwIL-15 variants. (Right) Schematic depiction of the various selected linker combinations. **c-d**Pre-activated mouse T cells were induced with 5 nM MMP-L_1_-SwIL-15 (**c**), MMP-L_2_-SwIL-15 (**d**) with or without pre-treatment of 2 μg/mL MMP-2. **e**, MMP-2-dependent dose-response in mouse T cells induced with 5 nM MMP-responsive cytokines and MMP-2 (**f**). Throughout, data represent the mean ± s.e.m.

## Discussion

Cytokine-based therapies have emerged as promising immunotherapeutic approaches, offering powerful tools to modulate the immune response. However, the clinical application of cytokines has been limited by their pleiotropic effects and associated toxicities^38–42^. To address this challenge, the concept of leveraging distinct tumor properties to regulate cytokine activity conditionally holds great potential. Previous efforts have leveraged tumor-intrinsic characteristics^4–6,11,12,17,29^, thus allowing conditional cytokine activation in the TME, but lacking precise control over the timing of cytokine activity, a critical factor in clinical settings.

Herein, we presented a novel strategy for achieving precise control of cytokine activity using the clinically approved drug Venetoclax. This approach involves the fusion of Bcl-2 to the cytokine of interest and incorporates the BIM-BH3 interaction motif at sites proximal to the cytokine’s receptor binding site. Using this design, we engineered switchable variants of both monomeric cytokines, such as IL-2 and IL-15, and structurally more complex cytokines, such as the dimeric IL-10. Employing IL-15 as an example cytokine, we demonstrated that in the absence of Venetoclax, Bcl-2 bound to the cytokine with high affinity, thus efficiently blocking the interaction site between the cytokine and its receptor. Upon drug administration, Venetoclax successfully disrupted the BIM-Bcl-2 interaction, hence restoring cytokine activity *in vitro* and *in vivo*.

Fc-fused cytokines are used in patients for increased half-life, allowing them to persist longer following a single infusion. While this prolonged exposure is advantageous by enabling less frequent dosing, it may also trigger adverse effects. The SwIL platform offers a means to engineer cytokine therapeutics with more precise dosing and potentially improved safety profiles. Furthermore, SwIL-based therapeutics with temporal control may facilitate step-up dosing strategies and enable rapid reversibility of therapeutic activity to alleviate chronic stimulation of target T cells and prevent T-cell exhaustion, thereby enhancing therapeutic efficacy^43,44^.

Switching kinetics are a critical consideration in the design of stimulus-responsive molecules. In this study, we demonstrated that our strategy enables fine-tuning of switching kinetics by modulating the intramolecular interaction between Bcl-2 and the BIM peptide. Functional assays showed full activation of SwIL molecules in the range of minutes upon incubation with the drug.

Previous efforts to engineer conditional cytokine activity relied on the fusion of one domain of the cytokine’s receptor to the cytokine via an MMP-cleavable linker^12,13,17,18^. While these strategies proved effective, the employment of the receptor’s subunit could limit the generalizability of the strategy across different cytokines. In this study, we showed that our engineering approach imparts switchable activity to monomeric (IL-2 and IL-15) and dimeric (IL-10) proteins, indicating the potential applicability of this strategy to a broader range of cytokines. While drug responsiveness offers significant advantages, achieving tumor-specific activation of these proteins would further enhance their therapeutic potential. To explore this, we demonstrated the modularity of our system by incorporating MMP-cleavable linkers between the SwIL and Bcl-2. All designs tested exhibited MMP-induced activation to varying extents. Additionally, a dose-response analysis of three designs showed variable activation profiles in response to MMP concentrations, suggesting the potential to tune the activation of this system in response to MMP.

Immunogenicity is a critical factor to consider in the design of switchable therapeutics. The SwIL system was constructed entirely from human protein components, and we therefore speculate that the human origin of these sequences may reduce the risk of immunogenic responses. Moreover, a current limitation of the proposed SwIL system is the helical nature of the BIM peptide, which will likely only be structurally compatible with cytokines with helical structures.

The engineered cytokines demonstrated switchability *in vivo* in mouse tumor models, paving the way for future clinical applications. Survival of tumor-bearing mice was substantially prolonged in those treated with SwlL-15 in combination with Venetoclax, achieving survival rates comparable to those of the unmodified cytokine. In the selected model, the combined treatment with ACT of antigen-specific T cells and IL-15/Fc did not lead to any severe toxicities. However, in the clinic, many promising cytokines, such as IL-15 superagonist (IL-15SA) or IL-12, often exhibited potent dose-limiting toxicities when combined with other immunotherapies^45,46^. Therefore, our strategy attempts to enhance the safety profile of these cytokines for the treatment of cancer patients, opening possibilities for their integration into antigen-targeted immunocytokines or armored CAR-T cell constructs.

Smart biologics with conditional activity offer promising strategies to enhance both efficacy and safety. Here, we developed a switchable cytokine system responsive to both drug and tumor-associated stimuli. Our results demonstrate that the system can be implemented across cytokines with diverse structural folds, highlighting the generalizability of the approach and advancing the development of smart biologics with translational potential.

## Methods

### Animals, cell lines, and reagents

All the mouse studies were approved by the Swiss authorities (Canton of Vaud, animal protocol ID 3902) and performed by guidelines from the Center of PhenoGenomics (CPG) in EPFL. 6-to 8-week-old female C57BL/6 mice were purchased from Charles River Laboratories (Lyon, France). T-cell receptor (TCR)-transgenic Thy1.1^+^Pmel-1 (Pmel) mice (B6.Cg-*Thy1*a/cyTg(TcraZcrb)8Rest/J) were purchased from The Jackson Laboratory (Bar Harbor, ME, USA) and maintained in the animal facility in the CPG in EPFL.

B16F10 murine melanoma cells were originally acquired from the American Type Culture Collection (ATCC; Manassas, VA, USA). HEK-Blue^™^IL-2 cells were originally acquired from Invivogen (San Diego, CA, USA). HEK293T cells were originally acquired from Invitrogen (Ref: R70007) and Expi293TM cells were originally acquired from ThermoFisher (Ref: A14635). B16F10, HEK-Blue^™^IL-2 and HEK293T cells were cultured in Dulbecco’s modified Eagle’s medium (DMEM) (Gibco, Thermo Fisher Scientific, Waltham, MA, USA) supplemented with fetal bovine serum (FBS) (10 v/v%, Gibco) and penicillin/streptomycin (1 v/v%, Gibco).

4′,6-diamidino-2-phenylindole dihydrochloride (DAPI) was purchased from Sigma-Aldrich (St. Louis, MO, USA). Recombinant human interleukin-2 (IL-2), interleukin-15 (IL-15), and interleukin-10 (IL-10) were purchased from PeproTech (London, UK). The luciferase detection kit was purchased from Promega (N1110, Madison, WI, USA). Venetoclax (ABT-199) was purchased from ChemieTek (Indianapolis, IN, USA). Polyethyleneimine (PEI) was purchased from Polysciences (Warrington, PA, USA).

For flow cytometry analysis, the following antibodies were purchased from BD Biosciences (Franklin Lakes, NJ, USA): anti-STAT5 (pY694) (47, 612599) and anti-pSTAT3 (4, 557815).

### Design of SwILs

The BIM motif sequence (IAQELRRIGDEF) was added onto the N or C-termini of interleukins. Given the high affinity of the superkine to IL-2R, we incorporated an extended version of the BIM peptide (DMRPEIWIAQELRRIGDEF) into the SwIL-2 variants. Design variants included deletions of non-receptor-binding terminal interleukin residues or the addition of alanine residues between the interleukin and BIM motif, generating variants that differ in their orientation of the bound Bcl2. Flexible (GS)_n_linkers were incorporated between the BIM and the previously optimized version of Bcl-2^47^. SwIL variants were modelled using ColabFold, with MMseqs2 used for MSAs. The output interleukin segments were superimposed, using PyMol 2.0, to existing structures of each interleukin bound to their respective receptor (PDB: IL2 - 2ERJ, IL10 - 6X93, IL15 – 4GS7). Models that showed clashes between their bound Bcl-2 placement and the receptor were selected for screening. Protein sequences are listed in tables S1 – S5.

### Computational SSM

The Rosetta protein modeling suite was utilized to identify BIM mutants that destabilize its interface with Bcl-2, thereby facilitating faster activation upon the addition of Venetoclax. To calculate the binding energy for the intramolecular interaction between the single-chain SwIL-15’s BIM and Bcl-2, first, the pose was relaxed using Rosetta’s main ‘Relax’ protocol. The linker was removed from the model, and the Interleukin+BIM and Bcl-2 were split into two separate chains. The site-saturation mutagenesis was performed using a RosettaScripts protocol, which iteratively selects each residue on the BIM peptide and mutates it to every other canonical amino acid using the Task Operation ‘RestrictAbsentCanonicalAASRLT’ while preventing all other residues in the pose from being repacked using the Task Operation ‘PreventRepackingRLT.’ Each mutation was incorporated into the pose using the Mover ‘PackRotamersMover’. For each mutation, scores were generated using the Filters ‘EnergyPerResidue’ (EPR) to evaluate the local stability and ‘Ddg’ to assess the mutation’s effect on binding affinity. Three alanine mutants were selected for characterization to conserve the helicity of the BIM peptide and the integrity of the interface. These mutants deviated from their respective native BIM residues with a change in REUs of <2.5 for the EPR, two had an REU change in Ddg of <5, while one was selected to have an REU change in Ddg of <10.

### Design of MMP-responsive SwIL variants

Three different MMP cleavage sites were flanked by different combinations of linkers and integrated into the SwIL-15_mut11 variant between the BIM and Bcl-2 regions. These designs consisted of MMP cleavage sites L1: (HPVGLLAR)_3_, L2: (VPLSLYSG)_3_, and L3: (HPVGLLAR)_2_(VPLSLYSG). These MMP sites incorporated either a flexible (GS)_n_or rigid linker (EAAAK or PAPAP) at each side. Designs were modelled using ColabFold with MMseq2 used for MSAs. Predicted structures of designs were manually analyzed to ensure the BIM:Bcl-2 interface was conserved, and the MMP cleavage site was adequately exposed. Genes for eight designs were ordered, expressed, and purified for further analysis.

### Protein expression and purification

For the production of wild-type (WT) cytokines, optimized sequences encoding human IL-10, IL-15, and superkine were synthesized and fused to a non-cytolytic human IgG1 Fc. A sequence encoding a (His)_6_-tag was fused to the C terminus of all proteins. DNA sequences were ordered from Twist Bioscience (South San Francisco, CA, USA) and cloned into the pCDNA mammalian expression vector through Gibson cloning. Mammalian expressions were performed using the Expi293^™^expression system from ThermoFisher Scientific (Waltham, MA, USA). The supernatant was collected 6 days post-transfection, filtered, and purified. Proteins were then purified using an ÄKTA pure system (GE Healthcare, Chicago IL, USA) with Ni-NTA affinity columns followed by size exclusion chromatography with PBS. The purified proteins were aliquoted and stored at −80 °C before use.

### HEK-293T transfection

For expression of cytokines or reporter system in human embryonic kidney cells (HEK-293T, ATCC: CRL-11268), 1 ×10^4^/well were seeded into 96-well plates 24 hours before transfection. The transfection mix in each well consisted of 125-130 ng of plasmid DNA mixed with 50 μL of FBS and antibiotic-free media and 600 ng PEI. The transfection mix was prepared, thoroughly mixed, and incubated at room temperature for 20-30 min. Then, it was added to the cells for overnight transfection. The next day, the medium was exchanged with fresh pre-warmed complete culture medium containing the appropriate inducer concentration or no inducer.

### IL-2 and IL-15 binding assays in HEK-Blue^™^cells

HEK-Blue^™^IL-2 cells were seeded into 96-well plates at a density of 1 ×10^4^cells/well 24 hours before treatment. In the morning, the medium was exchanged with 100 μL of medium containing different concentrations of cytokine with or without drug or no cytokine as a negative control. SEAP (human placental-secreted alkaline phosphatase) output was measured 24 hours after induction for all experiments.

### Reporter measurements in HEK-293T cells

SEAP concentrations in cell culture supernatants were quantified in terms of absorbance increase due to hydrolysis of para-nitrophenyl phosphate (pNPP). 80 μL of heat-inactivated (30 min at 65 °C) supernatants was mixed in a 96-well plate with 100 μL of 2x SEAP buffer (20 mM homoarginine, 1 mM MgCl_2_, 21% (v/v) diethanolamine, pH 9.8) and 20 μL of substrate solution containing 20 mM pNPP (Acros Organics BVBA). Samples were measured at 405 nm every minute for 30 min at 37°C on a Tecan Spark plate reader (Tecan AG). NanoLuc concentrations in cell culture supernatants were quantified with the Nano-Glo Luciferase Assay System (N1119, Promega). 5 μL of the sample were incubated for 5 min in a 384-well plate with 5 μL substrate/buffer mix (1:50). Luminescence was measured on a Tecan Spark plate reader (Tecan AG).

### Culture of primary T cells

Spleens from C57BL/6 or Pmel mice were mechanically disrupted into individual cells by smashing on a 70-μm strainer (Fisher Scientific, Pittsburgh, PA, USA) and then lysed with ACK Lysis buffer (1 mL per spleen, Gibco/Thermo Fisher Scientific) for 5 min to remove red blood cells. Splenocytes were then washed with PBS and resuspended to a cell density of around 1.0 × 10^6^/mL with complete RPMI 1640 culture medium (Gibco/Thermo Fisher Scientific) containing FBS (10%, v/v), HEPES (1%, v/v), penicillin/streptomycin (1%, v/v), and β-mercaptoethanol (0.1%, v/v) supplemented with mouse IL-2 (10 ng/mL; PeproTech, London, UK), IL-7 (1 ng/mL; PeproTech), and gp100 25–33 (1 μM; GenScript, Piscataway, NJ, USA). 3 days after activation, live cells were enriched by density gradient centrifugation with Ficoll-Paque PLUS (GE Healthcare). The collected cells were cultured for an extra 2 days at a cell density of around 1.0 × 10^6^/ml in complete RPMI 1640 medium supplemented with mouse IL-2 (10 ng/mL) and IL-7 (2 ng/mL). WT mouse T cells were purified from spleens of C57BL/6 mice using mouse Pan T cell isolation kit (Miltenyi Biotec, Bergisch Gladbach, DE). Purified CD3^+^T cells (10 × 10^6^cells/well) were activated in 6-well plates pre-coated with 1 μg/mL anti-CD3 (clone 17A2, BioxCell) and supplemented with 5 μg/mL anti-CD28 (37.51, BioXCell) and 10 ng/mL mouse IL-2 (Peprotech) for 3 days. After 3 days of activation, T cells were detached from the activating antibodies and continued culturing at a cell density of 0.5-1 x 10^6^cells/mL in 10 ng/mL mIL-2. Human peripheral blood mononuclear cells (PBMCs) of healthy donors (prepared as a buffy coat) were separated by Ficoll-Paque PLUS density gradient media (Cytiva, Marlborough, MA, USA) centrifugation. Human T cells were isolated using the human Pan T cell Isolation Kit (130-096-535, Miltenyi Biotec, Bergisch Gladbach, DE) and stimulated with anti-CD3 and ani-CD28 monoclonal antibody (mAb)-coated beads (Invitrogen, Life Technologies) at a ratio of 1:2 T cells to beads and 50 U/mL recombinant human IL-2 (Peprotech, London, UK). Human IL-2 was replenished every other day at a concentration of 50 U/mL. A cell density of 0.5-1 x 10^6^cells/mL was maintained for expansion.

### Analysis of STAT5 phosphorylation by flow cytometry

Activated T cells were starved overnight in complete culture medium without cytokines. The next day, starved cells were first cultured in media lacking fetal bovine serum (FBS) for 2 hours at 37 °C and 5% CO_2_. Next, cells were plated in pre-warmed complete culture medium (1 × 10^5^cells/well) and indicated amounts of WT cytokine (IL-2, IL-10/Fc, or IL-15/Fc) or switchable cytokine (SwIL-2, SwIL-10, or SwIL-15) were applied to T cells for 15-18 min at 37°C to induce STAT5 phosphorylation. Cells were fixed immediately with 2% paraformaldehyde for 10 min at room temperature anti-pSTAT5 antibodies (1:50 dilution, BD Biosciences) for 2 hours at room temperature in the dark. Cells were acquired using Attune NxT Flow Cytometer with Attune NxT Software v.3 (Invitrogen/ThermoFisher Scientific). Data analysis was performed using FlowJo 10.6.1 (Tree Star).

### Switching kinetics analysis

5 nM of cytokine were incubated with and without 10 μM Venetoclax for 0.3, 1, 2, and 16 hours at 37°C. The mixture was then used to induce STAT5 phosphorylation, as described above.

### Cleavage of SwIL-15 by recombinant proteases

Active recombinant human MMP-2 protein (ab283047) was purchased from Abcam. Recombinant active MMP-2 and SwIL-15 were diluted in a TNBC buffer containing 50 mM Tris-HCl, pH 7.5, 150 mM NaCl, 5 mM CaCl_2_, 0.025% Brij-35, and incubated at 37°C for 1 hour. MMP-2-treated cytokines were then used for pSTAT5 stimulation in pre-activated CD8^+^T cells. Final concentrations of MMP-2 and cytokines are indicated in the figure legends.

### Tumor models

B16F10 cells (0.5 × 10^6^) in PBS (100 μL) were inoculated subcutaneously in the right flanks of female C57BL/6J mice at day 0. Mice bearing palpable tumors between 15-35 mm^2^were randomized on day 6 post-inoculation and were lymphodepleted at 4 Gy irradiation, and adoptive cell transfer (ACT) of activated Pmel T cells (5 × 10^6^cells) was performed via intravenous (i.v.) injection at day 7. Tumor area (product of measured orthogonal length and width) and body weight were monitored every 2 days. Mice were euthanized when the body weight loss was higher than 20% of the pre-dosing weight or the tumor area reached 150 mm^2^. Mice bearing B16F10 tumors received intratumoral (i.t.) injections of IL-15/Fc (200 pmol/dose in 50 μL PBS) or SwIL-15 (200 pmol/dose in 50 μL PBS) every second day for a total of 5 doses, and daily subcutaneous (s.c.) injections of Venetoclax (25 mg/kg in 50 μL PBS) from day 7 to 15. Mice receiving PBS, Venetoclax, or IL-15/Fc served as controls.

### *In vivo*drug toxicity testing

Healthy C57BL/6 female mice, aged 6-10 weeks, were shaved in the right flank and treated daily with 50 μL s.c. injections of Venetoclax dissolved at 25 mg/kg in PBS and 2% dimethyl sulfoxide (DMSO) or vehicle (2% DMSO). The animals were monitored every second day to assess any sign of drug toxicity evaluated by body weight loss.

### Pharmacokinetic study of SwIL-15

Healthy C57BL/6 female mice received equimolar treatment (100 pmol) of IL-15/Fc or SwIL-15 intraperitoneally (i.p.), and mice (n=3 per group) were bled 0.5, 1, 2, 4, 8, 24, and 48 hours post-injection. Plasma was separated and analyzed by ELISA. ELISA plates were coated with anti-6x-His Tag antibody (HIS.H8, MA1-21315, Invitrogen) overnight at 4°C. Following, plasma was added to the wells and incubated, and the presence of hIL-15/Fc or SwIL-15SA was detected using a horseradish peroxidase (HRP)-conjugated anti-human IgG antibody. hIL-15/Fc and SwIL-15 generated in our lab were used as standards. Plasma half-life was estimated using a one-phase decay model (v.10, GraphPad Prism).

### Statistical analysis

Data are presented as mean ± standard error of the mean (SEM) unless otherwise noted. Statistical analysis for each experiment is specified in the corresponding figure legend. Statistical analyses were performed using GraphPad Prism 8 software. Two-tailed tests with P values of less than 0.05 were considered significant in all cases.

## Acknowledgments

We acknowledge the EPFL Center of PhenoGenomics, Flow Cytometry Core Facility, and Protein Expression Core Facility for technical assistance. B.E.C. was supported by the Swiss National Science Foundation, the NCCR in Chemical Biology, and the NCCR in Molecular Systems Engineering. L.T. acknowledges the grant support from the Swiss National Science Foundation (grant nos. 315230_204202, IZLCZ0_206035, CRSII5_205930), European Research Council under European Research Council grant agreement MechanoIMM (grant no. 805337), Swiss Cancer Research Foundation (grant no. KFS-4600-08-2018), and EPFL.

## Author contributions

L.B. and S.B. contributed equally to this work. L.B., L.S., S.S., L.T., and B.E.C. conceived the project. L.B., S.B., L.T., and B.E.C. designed experiments and experimental methodology. L.B., S.B., L.S., A.M., and T.E. performed the experimental work. L.B. and S.B. performed the computational work and protein design. S.G. and S.B. performed the expression and purification of proteins. L.T. and B.E.C. provided supervision and acquired the necessary funding. L.B., S.B., L.T., and B.E.C. wrote the manuscript with input from all authors.

