## Supplementary Information for "Rational design of chemically responsive cytokines for cancer immunotherapy"

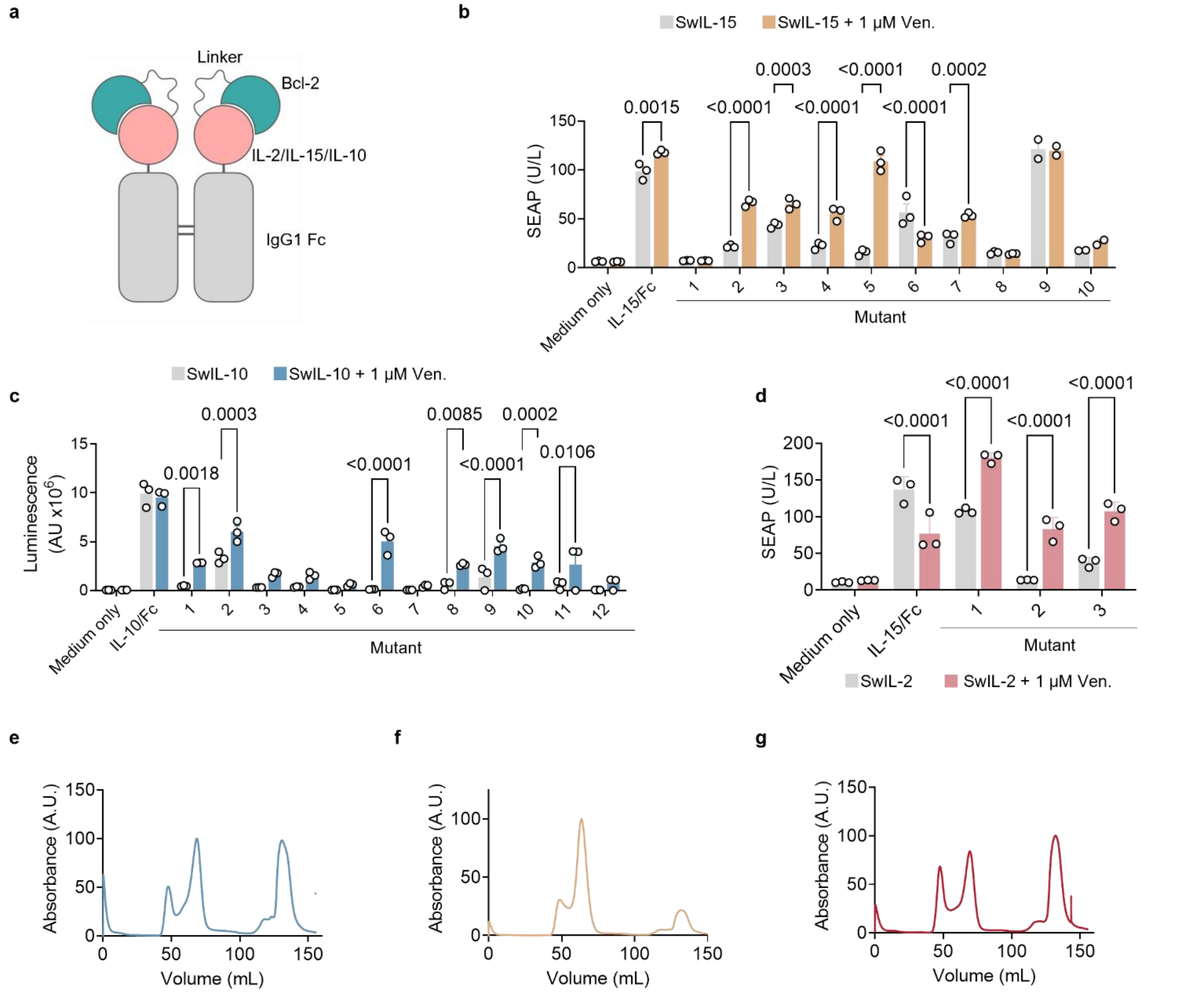

h

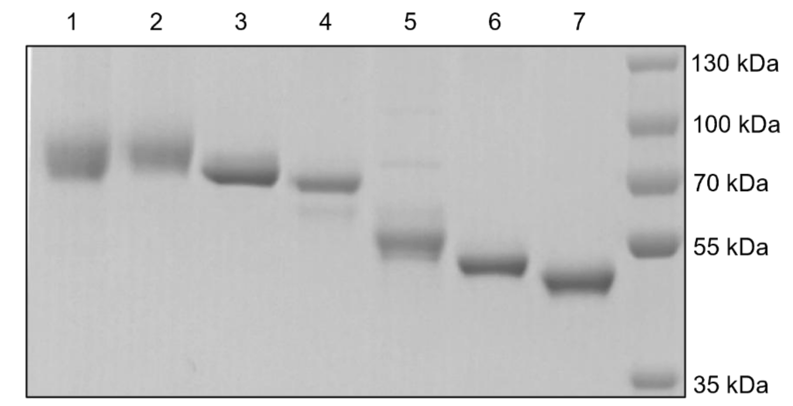

| Line | Protein | Expected size (kDa) |
|---|---|---|
| 1 | SwIL-15_mut11 | 61.9 |
| 2 | SwIL-15_mut5 | 61.7 |
| 3 | SwIL-10_mut10 | 70.0 |
| 4 | SwIL-2_mut2 | 63.4 |
| 5 | hIL-15/Fc | 40.3 |
| 6 | hIL-10/Fc | 48.3 |
| 7 | Superkine/Fc | 41.3 |

**Supplementary Fig. 1 Screening of switchable cytokine mutants and size-exclusion chromatograms of affinity-purified switchable cytokines. a**, Schematic diagram of switchable cytokine constructs. **b-d**, Screening of switchable cytokine mutants in HEK293T cells of SwIL-15 (**b**), SwIL-10 (**c**), and SwIL-2 (**d**). **e-g**, $(His)_6$-tagged SwIL-15 Mut5 (**e**), SwIL-10 Mut10 (**f**), and SwIL-2 Mut2 (**g**) were expressed in Expi293™ cells and purified via Nickel-based affinity purification as described in the Methods section. **h**, Representative sodium dodecyl sulfate-polyacrylamide gel electrophoresis (SDS-PAGE) analysis of purified SwIL-15_mut11, SwIL-15_mut5, SwIL-10_mut10, SwIL-2_mut2, hIL-15/Fc, hIL-10/Fc, superkine/Fc. P-values were determined by unpaired Student's t-test. (**b**-**d**).

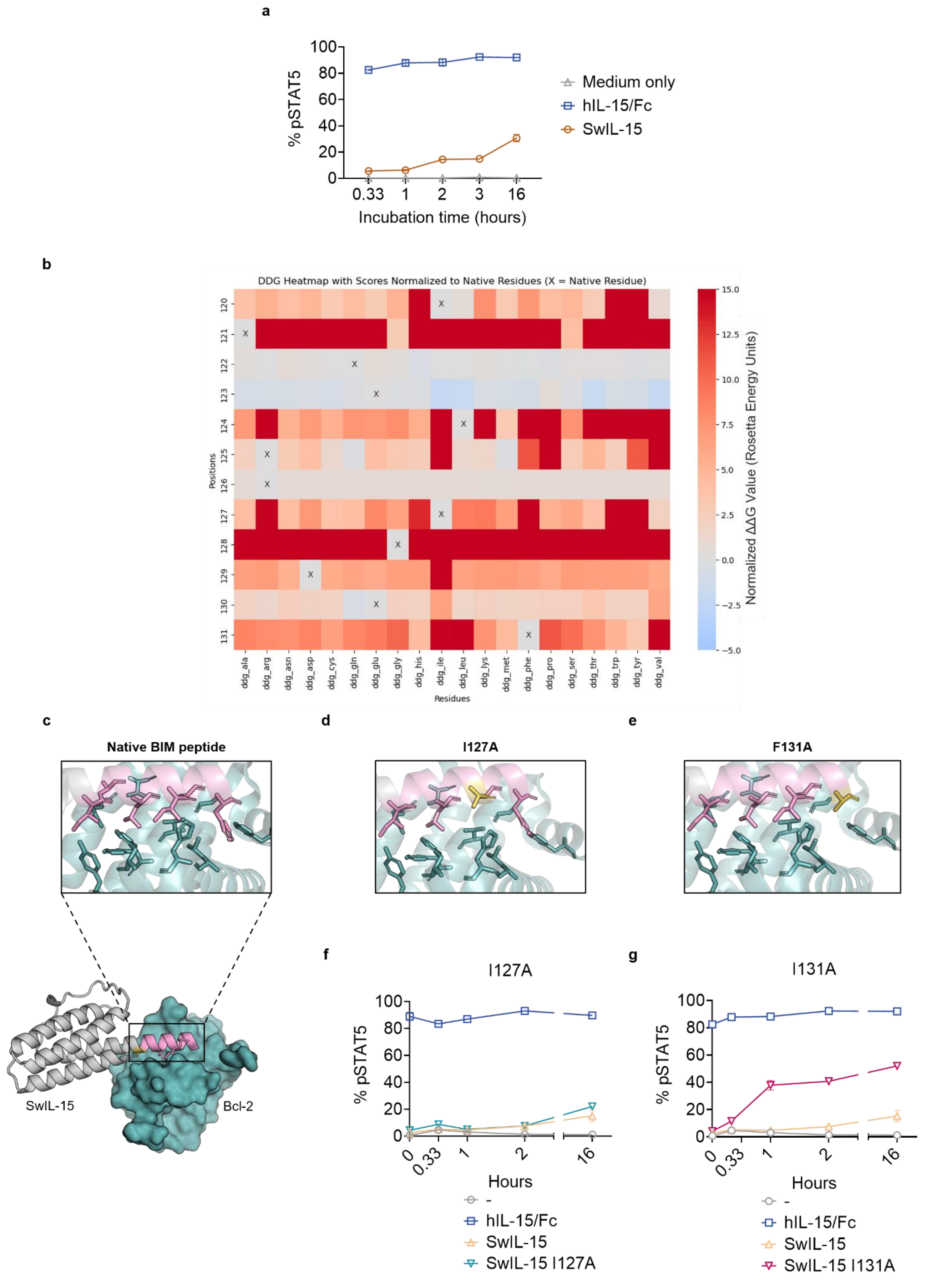
a
100
80
60
40
20
0
% pSTAT5
0.33
1
2
3
16
Incubation time (hours)
Medium only
hIL-15/Fc
SwIL-15
b
DDG Heatmap with Scores Normalized to Native Residues (X = Native Residue)
Positions
Residues
ddg_ala
ddg_arg
ddg_asn
ddg_asp
ddg_cys
ddg_gln
ddg_glu
ddg_gly
ddg_his
ddg_ile
ddg_leu
ddg_lys
ddg_met
ddg_phe
ddg_pro
ddg_ser
ddg_thr
ddg_trp
ddg_tyr
ddg_val
Normalized ΔΔG Value (Rosetta Energy Units)
15.0
12.5
10.0
7.5
5.0
2.5
0.0
−2.5
−5.0
c
Native BIM peptide
SwIL-15
Bcl-2
d
I127A
e
F131A
f
I127A
% pSTAT5
Hours
0
0.33
1
2
16
-
hIL-15/Fc
SwIL-15
SwIL-15 I127A
g
I131A
% pSTAT5
Hours
0
0.33
1
2
16
-
hIL-15/Fc
SwIL-15
SwIL-15 I131A

**Supplementary Fig. 2 Tuning the affinity of Bcl-2 to the BIM peptides results in differential switching kinetics.** **a**, Switching kinetics of SwIL-15. 5 nM of SwIL-15 or IL-15/Fc were incubated with 10 µM Venetoclax for 0.33, 1, 2, 3, and 16 hours at 37°C and then employed to induce STAT5 phosphorylation in primary mouse T cells. **b**, Computational site saturation mutagenesis obtained with Rosetta. Mutations to amino acids resulting in an increase of the computed binding energy (ΔΔG) of at least two Rosetta energy units (REU) were considered as mutant candidates. **c**, Model of SwIL-15 interacting with Bcl-2. **d-e**, Models of the BIM peptide mutations I127A (**d**) and F131A (**e**). **f-g**, Switching kinetics of SwIL-15 I127A (**f**) and F131A (**g**).

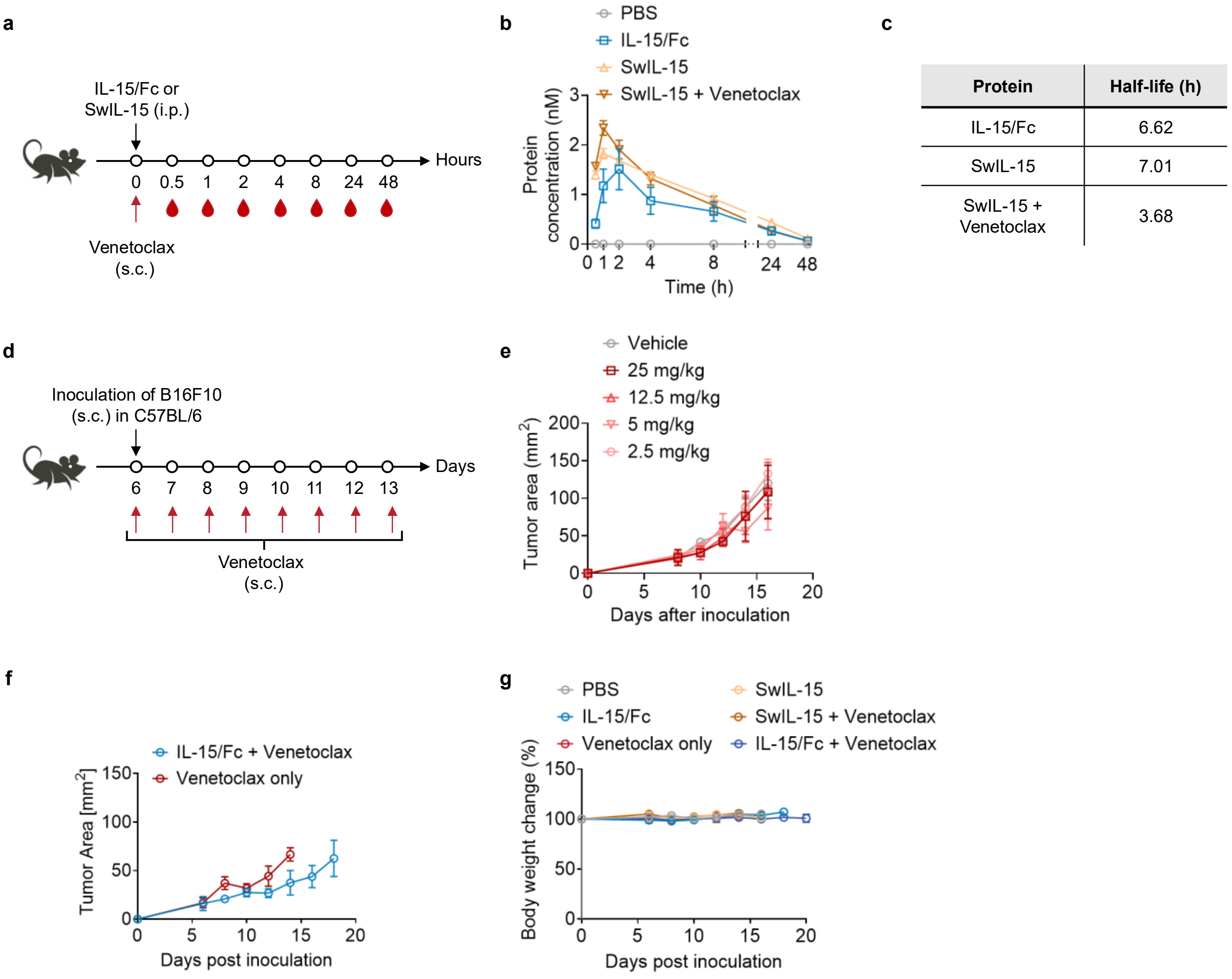

| Protein | Half-life (h) |
|---|---|
| IL-15/Fc | 6.62 |
| SwIL-15 | 7.01 |
| SwIL-15 + Venetoclax | 3.68 |

**Supplementary Fig. 3 SwIL-15 shows half-life in the same range of is unmodified counterpart and Venetoclax does not affect tumor growth nor IL-15 performance. a**, Healthy C57BL76 mice treated i.p. with 100 pmol of IL-15/Fc, SwIL-15, or vehicle with or without 25 mg/kg Venetoclax (injected s.c.) and bled at the indicated time points (n = 3 animals per group). Plasma was analyzed for IL-15 concentration via ELISA. **b-c**, Shown are half-life decay curves (**b**) and half-time values (**c**). **d**, C57BL/6 mice were s.c. injected with B16F10 melanoma tumor cells (2 × $10^5$ cells) and starting from day 6 received daily s.c. injections of Venetoclax at the indicated concentrations. **e**, Shown is the tumor growth curve. No significant difference was observed between the groups. **f**, C57BL/6 mice were inoculated subcutaneously (s.c.) with B16F10 melanoma tumor cells (3 × $10^5$), lymphodepleted by X-ray irradiation (4 Gy), and received adoptive transfer of $CD8^+$ T cells (5 × $10^6$) on day 6. Tumor-bearing mice were treated with IL-15/Fc (100 pmol/injection) and or SwIL-15 (100 pmol/injection) every other day with or without daily Venetoclax administration (25 mg/kg). **f-g**, Shown are tumor growth curves of mice treated with IL-15 + Venetoclax or Venetoclax only (**f**), and body weight change of all groups (**g**).

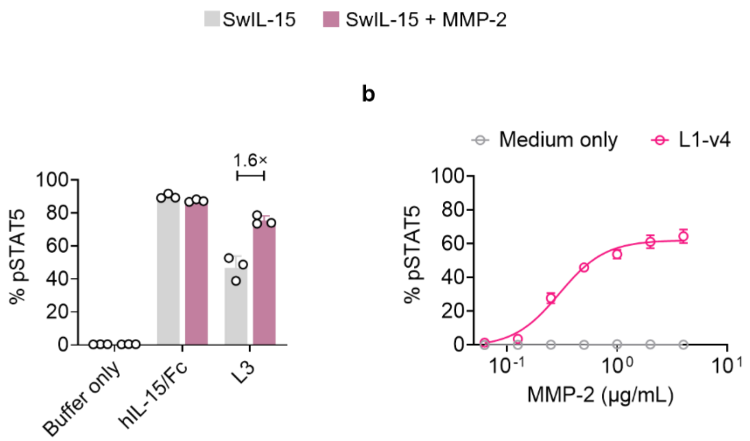

**Supplementary Fig. 4 MMP-responsive SwIL-15 show dose-responsive behavior. a**, Pre-activated mouse primary T cells were induced with 5 nM MMP-$L_3$-SwIL-15 with or without pre-treatment with 2µg/mL MMP-2. MFI: Mean fluorescence intensity. **b**, MMP2-dependent dose-response of phosphorylated STAT5 (pY694) with MMP2-treated MMP-$L_1$-v4-SwIL-15 mutants in pre-activated primary mouse T cells. MFI, mean fluorescence intensity

**Supplementary Table S1: Bcl-2 sequence.** Sequence of the optimized Bcl-2 protein used in SwILs.

| **Bcl-2 sequence** | | |
|---|---|---|
| **Protein** | **Sequence** | **Description** |
| Bcl-2 | AHAGRTGYDNREIVMKYIHYKLSQRGYEWDAGDDAEENRTEAPEGTESEVVHRALRDAGDDFERRYRRDFAEMSSQLHLTPDTARQRFETVVEELFRDGVNWGRIVAFFEFGGVMCVESVNREMSPLVDNIAEWMTEYLNRHLHTWIQDNGGWDAFVELYGPSMRGGGGS | Optimized Bcl-2 |

**Supplementary Table S2: SwIL-15 designs.** List of 10 screened SwIL-15 designs. Underlined: BIM peptide. Bold: residue additions.

| SwIL-15 designs | | |
|---|---|---|
| Design | Sequence | Description |
| SwIL-15 Mutant 1 | IAQELRRIGDEFAAAMAISNWVNVISDLKKIEDLIQSMHIDATLYTESDVHPSCKVTAMKCFLLELQVISLESGDASIHDTVENLIILANNSLSSNGNVTESGCKECEELEEKNIKEFLQSFVHIVQMFINTSASGGGGSGGGGSGGGGS[Bcl-2] | N-terminal BIM + 2x Ala addition |
| SwIL-15 Mutant 2 | IAQELRRIGDEFAAAAMAISNWVNVISDLKKIEDLIQSMHIDATLYTESDVHPSCKVTAMKCFLLELQVISLESGDASIHDTVENLIILANNSLSSNGNVTESGCKECEELEEKNIKEFLQSFVHIVQMFINTSASGGGGSGGGGSGGGGS[Bcl-2] | N-terminal BIM + 3x Ala addition |
| SwIL-15 Mutant 3 | IAQELRRIGDEFAAAAAMAISNWVNVISDLKKIEDLIQSMHIDATLYTESDVHPSCKVTAMKCFLLELQVISLESGDASIHDTVENLIILANNSLSSNGNVTESGCKECEELEEKNIKEFLQSFVHIVQMFINTSASGGGGSGGGGSGGGGS[Bcl-2] | N-terminal BIM + 4x Ala addition |

| | | |
|---|---|---|
| SwIL-15 Mutant 4 | IAQELRRIGDEFAAAAAAAMAISNWVNVISDLKKIEDLIQSMHIDATLYTESDVHPSCKVTAMKCFLLELQVISLESGDASIHDTVENLIILANNSLSSNGNVTESGCKECEELEEKNIKEFLQSFVHIVQMFINTSASGGGGSGGGGSGGGGS[Bcl-2] | N-terminal BIM + 6x Ala addition |
| SwIL-15 Mutant 5 | AMAISNWVNVISDLKKIEDLIQSMHIDATLYTESDVHPSCKVTAMKCFLLELQVISLESGDASIHDTVENLIILANNSLSSNGNVTESGCKECEELEEKNIKEFLQSFVHIVQMFINTSIAQELRRIGDEFASGGGGSGGGGSGGGGS[Bcl-2] | C-terminal BIM |
| SwIL-15 Mutant 6 | AMAISNWVNVISDLKKIEDLIQSMHIDATLYTESDVHPSCKVTAMKCFLLELQVISLESGDASIHDTVENLIILANNSLSSNGNVTESGCKECEELEEKNIKEFLQSFVHIVQMFINTSAAAIAQELRRIGDEFASGGGGSGGGGSGGGGS[Bcl-2] | C-terminal BIM + 3x Ala |
| SwIL-15 Mutant 7 | AMAISNWVNVISDLKKIEDLIQSMHIDATLYTESDVHPSCKVTAMKCFLLELQVISLESGDASIHDTVENLIILANNSLSSNGNVTESGCKECEELEEKNIKEFLQSFVHIVQMFINTSAAAAIAQELRRIGDEFASGGGGSGGGGSGGGGS[Bcl-2] | C-terminal BIM + 4x Ala |
| SwIL-15 Mutant 8 | AMAISNWVNVISDLKKIEDLIQSMHIDATLYTESDVHPSCKVTAMKCFLLELQVISLESGDASIHDTVENLIILANNSLSSNGNVTESGCKECEELEEKNIKEFLQSFVHIVQMFINTSAAAAIAQELRRIGDEFNAYYARASGGGGSGGGGSGGGGS[Bcl-2] | C-terminal full-length BIM + 4x Ala |
| SwIL-15 Mutant 9 | AMAISNWVNVISDLKKIEDLIQSMHIDATLYTESDVHPSCKVTAMKCFLLELQVISLESGDASIHDTVENLIILANNSLSSNGNVTESGCKECEELEEKNIKEFLQSFVHIVQMFINTSDALKIAQELRRIGDEFASGGGGSGGGGSGGGGS[Bcl-2] | C-terminal BIM + DALK |

| | | |
|---|---|---|
| SwIL-15 Mutant 10 | IAQELRRIGDEFNAYYAMAISNWVNVISDLKKIEDLIQSMHIDATLYTESDVHPSCKVTAMKCFLLELQVISLESGDASIHDTVENLIILANNSLSSNGNVTESGCKECEELEEKNIKEFLQSFVHIVQMFINTSASGGGGSGGGGSGGGGS[Bcl-2] | N-terminal BIM + NAYY |

**Supplementary Table S3: SwIL-10 designs.** List of 10 screened SwIL-15 designs. Underlined: BIM peptide. Bold: residue additions.

| SwIL-10 designs | | |
|---|---|---|
| Design | Sequence | Description |
| SwIL-10 Mutant 1 | SPGQGTQSENSCTHFPGNLPNMLRDLRDAFSRVKTFFQMKDQLDNLLLKESLLEDFKGYLGCQALSEMIQFYLEEVMPQAENQDPDIKAHVNSLGENLKTLRLRLRRCHRFLPCENKSKAVEQVKNAFNKLQEKGIYKAMSEFDIFINYIEAYMTMKIRNIAQELRRIGDEFGGSGGS[Bcl-2] | C-terminal BIM |
| SwIL-10 Mutant 2 | SPGQGTQSENSCTHFPGNLPNMLRDLRDAFSRVKTFFQMKDQLDNLLLKESLLEDFKGYLGCQALSEMIQFYLEEVMPQAENQDPDIKAHVNSLGENLKTLRLRLRRCHRFLPCENKSKAVEQVKNAFNKLQEKGIYKAMSEFDIFINYIEAYMTMKIRNGGSIAQELRRIGDEFGGSGGS[Bcl-2] | C-terminal BIM + $G_2S$ addition |
| SwIL-10 Mutant 3 | IAQELRRIGDEFSPGQGTQSENSCTHFPGNLPNMLRDLRDAFSRVKTFFQMKDQLDNLLLKESLLEDFKGYLGCQALSEMIQFYLEEVMPQAENQDPDIKAHVNSLGENLKTLRLRLRRCHRFLPCENKSKAVEQVKNAFNKLQEKGIYKAMSEFDIFINYIEAYMTMKIRNGGSGGS[Bcl-2] | N-terminal BIM |

| | | |
|---|---|---|
| SwIL-10 Mutant 4 | IAQELRRIGDEFGGSSPGQGTQSENSCTHFPGNLPNMLRDLRDAFSRVKTFFQMKDQLDNLLLKESLLEDFKGYLGCQALSEMIQFYLEEVMPQAENQDPDIKAHVNSLGENLKTLRLRLRRCHRFLPCENKSKAVEQVKNAFNKLQEKGIYKAMSEFDIFINYIEAYMTMKIRNGGSGGS[Bcl-2] | N-terminal BIM + $G_2S$ addition |
| SwIL-10 Mutant 5 | IAQELRRIGDEFCTHFPGNLPNMLRDLRDAFSRVKTFFQMKDQLDNLLLKESLLEDFKGYLGCQALSEMIQFYLEEVMPQAENQDPDIKAHVNSLGENLKTLRLRLRRCHRFLPCENKSKAVEQVKNAFNKLQEKGIYKAMSEFDIFINYIEAYMTMKIRNGGSGGS[Bcl-2] | N-terminal BIM + 11 residues truncation |
| SwIL-10 Mutant 6 | IAQELRRIGDEFGGSCTHFPGNLPNMLRDLRDAFSRVKTFFQMKDQLDNLLLKESLLEDFKGYLGCQALSEMIQFYLEEVMPQAENQDPDIKAHVNSLGENLKTLRLRLRRCHRFLPCENKSKAVEQVKNAFNKLQEKGIYKAMSEFDIFINYIEAYMTMKIRNGGSIAQELRRIGDEFGGSGGS[Bcl-2] | N- and C-terminal Bim + $G_2S$ addition at both termini + 11 residues truncation at the N-terminus |
| SwIL-10 Mutant 7 | SPGQGTQSENSCTHFPGNLPNMLRDLRDAFSRVKTFFQMKDQLDNLLLKESLLEDFKGYLGCQALSEMIQFYLEEVMPQAENQDPDIKAHVNSLGENLKTLRLRLRRCHRFLPCENKSKAVEQVKNAFNKLQEKGIYKAMSEFDIFINYIEAYMTMKIRNAAAAIAQELRRIGDEFGGSGGS[Bcl-2] | C-terminal BIM + AAAA |
| SwIL-10 Mutant 8 | SPGQGTQSENSCTHFPGNLPNMLRDLRDAFSRVKTFFQMKDQLDNLLLKESLLEDFKGYLGCQALSEMIQFYLEEVMPQAENQDPDIKAHVNSLGENLKTLRLRLRRCHRFLPCENKSKAVEQVKNAFNKLQEKGIYKAMSEFDIFINYIEAYMTMKIRNAAAIAQELRRIGDEFGGSGGS[Bcl-2] | C-terminal BIM + AAA |

| | | |
|---|---|---|
| SwIL-10 Mutant 9 | SPGQGTQSENSCTHFPGNLPNMLRDLRDAFSRVKTFFQMKDQLDNLLLKESLLEDFKGYLGCQALSEMIQFYLEEVMPQAENQDPDIKAHVNSLGENLKTLRLRLRRCHRFLPCENKSKAVEQVKNAFNKLQEKGIYKAMSEFDIFINYIEAYMTMKIRNAAIAQELRRIGDEFGGSGGS[Bcl-2] | C-terminal BIM + AA |
| SwIL-10 Mutant 10 | SPGQGTQSENSCTHFPGNLPNMLRDLRDAFSRVKTFFQMKDQLDNLLLKESLLEDFKGYLGCQALSEMIQFYLEEVMPQAENQDPDIKAHVNSLGENLKTLRLRLRRCHRFLPCENKSKAVEQVKNAFNKLQEKGIYKAMSEFDIFINYIEAYMTMKIRNAIAQELRRIGDEFGGSGGS[Bcl-2] | C-terminal BIM + A |
| SwIL-10 Mutant 11 | SPGQGTQSENSCTHFPGNLPNMLRDLRDAFSRVKTFFQMKDQLDNLLLKESLLEDFKGYLGCQALSEMIQFYLEEVMPQAENQDPDIKAHVNSLGENLKTLRLRLRRCHRFLPCENKSKAVEQVKNAFNKLQEKGIYKAMSEFDIFINYIEAYMTMKIAQELRRIGDEFGGSGGS[Bcl-2] | C-terminal BIM + 3 residues C-terminal truncation |
| SwIL-10 Mutant 12 | SPGQGTQSENSCTHFPGNLPNMLRDLRDAFSRVKTFFQMKDQLDNLLLKESLLEDFKGYLGCQALSEMIQFYLEEVMPQAENQDPDIKAHVNSLGENLKTLRLRLRRCHRFLPCENKSKAVEQVKNAFNKLQEKGIYKAMSEFDIFINYIEAYMTMKIRNIEEKIAQELRRIGDEFGGSGGS[Bcl-2] | C-terminal BIM + IEEK |
| SwIL-10 Mutant 13 | SPGQGTQSENSCTHFPGNLPNMLRDLRDAFSRVKTFFQMKDQLDNLLLKESLLEDFKGYLGCQALSEMIQFYLEEVMPQAENQDPDIKAHVNSLGENLKTLRLRLRRCHRFLPCENKSKAVEQVKNAFNKLQEKGIYKAMSEFDIFINYIEAYMTMKIRNDALKIAQELRRIGDEFGGSGGS[Bcl-2] | C-terminal BIM + DALK |

**Supplementary Table S4**: **List of 3 screened SwIL-2 designs.** Underlined: BIM peptide. Bold: mutations. Bcl-2 represents the sequence from Supplementary Table S1.

| SwIL-2 designs | | |
|---|---|---|
| Design | Sequence | Description |
| SwIL-2 Mutant 1 | DMRPEIWIAQELRRIGDEFKTQLQLEHLLLDLQMILNGINNYKNPKLTRMLTFKFYMPKKATELKHLQCLEEELKPLEEVLNLAQSKNFHLRPRDLISNINVIVLELKGSETTFMCEYADETATIVEFLNRWITFCQSIISTLTGGSGGS[Bcl-2] | N-terminal extended Bim.<br>Native hIL2 |
| SwIL-2 Mutant 2 | DMRPEIWIAQELRRIGDEFKTQLQLEHLLLDLQMILNMINNYDNPKLTRMLTFKFYMPKKATELKHLQCLEEELKPLEEVLNQAQSKNFHLDPRDLISNINVIVLELKGSETTFMCEYADETATIVEFLNRWITFCQSIISTLTGGSGGS[Bcl-2] | N-terminal extended Bim.<br>Superkine S15 rev |
| SwIL-2 Mutant 3 | DMRPEIWIAQELRRIGDEFKTQLQLEHLLLDLQMILNMINNYDNPKLTDMLTFEFYMPKKATELKHLQCLERELKPLEEVLNQAQSKNFHLDPRDLISNINVIVLELKGSETTFMCEYADETATIVEFLNRWITFCQSIISTLTGGSGGS[Bcl-2] | N-terminal extended Bim.<br>Superkine S15 rev + mutants |

**Supplementary Table S5**: **List of 8 screened MMP-responsive SwIL-15 designs.** Underlined: BIM peptide. Bold: mutations. Bcl-2 represents the sequence from Supplementary Table S1.

| MMP-SwIL-15 designs | | |
|---|---|---|
| Design | Sequence | Description |
| MMP-SwIL-15-L1-v1 | AMAISNWVNVISDLKKIEDLIQSMHIDATLYTESDVHPSCKVTAMKCFLLELQVISLESGDASIHDTVENLIILANNSLSSNGNVTESGCKECEELEEKNIKEFLQSFVHIVQMFINTSAAQELRRIGDEFASPAPAPHPVGLLARHPVGLLARHPVGLLARGGGSG[Bcl-2] | Rigid linker-$(L1)_3$-flexible linker |

| | | |
|---|---|---|
| MMP-SwIL-15-L1-v2 | AMAISNWVNVISDLKKIEDLIQSMHIDATLYTESDVHPSCKVTAMKCFLLELQVISLESGDASIHDTVENLIILANNSLSSNGNVTESGCKECEELEEKNIKEFLQSFVHIVQMFINTSAAQELRRIGDEFASEAAAKEAAAKHPVGLLARHPVGLLARHPVGLLARGGGSG[Bcl-2] | Rigid linker-(L1)$_3$-flexible linker |
| MMP-SwIL-15-L1-v3 | AMAISNWVNVISDLKKIEDLIQSMHIDATLYTESDVHPSCKVTAMKCFLLELQVISLESGDASIHDTVENLIILANNSLSSNGNVTESGCKECEELEEKNIKEFLQSFVHIVQMFINTSAAQELRRIGDEFASSGSGGGSHPVGLLARHPVGLLARHPVGLLARPAPAP[Bcl-2] | Flexible linker-(L1)$_3$-rigid linker |
| MMP-SwIL-15-L1-v4 | AMAISNWVNVISDLKKIEDLIQSMHIDATLYTESDVHPSCKVTAMKCFLLELQVISLESGDASIHDTVENLIILANNSLSSNGNVTESGCKECEELEEKNIKEFLQSFVHIVQMFINTSAAQELRRIGDEFASPAPAPHPVGLLARHPVGLLARHPVGLLARPAPAP[Bcl-2] | Rigid linker-(L1)$_3$-rigid linker |
| MMP-SwIL-15-L2-v1 | AMAISNWVNVISDLKKIEDLIQSMHIDATLYTESDVHPSCKVTAMKCFLLELQVISLESGDASIHDTVENLIILANNSLSSNGNVTESGCKECEELEEKNIKEFLQSFVHIVQMFINTSAAQELRRIGDEFASPAPAPVPLSLYSGVPLSLYSGVPLSLYSGGGGSG[Bcl-2] | Rigid linker-(L2)$_3$-flexible linker |
| MMP-SwIL-15-L2-v2 | AMAISNWVNVISDLKKIEDLIQSMHIDATLYTESDVHPSCKVTAMKCFLLELQVISLESGDASIHDTVENLIILANNSLSSNGNVTESGCKECEELEEKNIKEFLQSFVHIVQMFINTSAAQELRRIGDEFASEAAAKEAAAKVPLSLYSGVPLSLYSGVPLSLYSGGGGSG[Bcl-2] | Rigid linker-(L2)$_3$-flexible linker |
| MMP-SwIL-15-L2-v3 | AMAISNWVNVISDLKKIEDLIQSMHIDATLYTESDVHPSCKVTAMKCFLLELQVISLESGDASIHDTVENLIILANNSLSSNGNVTESGCKECEELEEKNIKEFLQSFVHIVQMFINTSAAQELRRIGDEFASGSGGGSVPLSLYSGVPLSLYSGVPLSLYSGPAPAP[Bcl-2] | Flexible linker-(L2)$_3$-rigid linker |
| MMP-SwIL-15-L3-v3 | AMAISNWVNVISDLKKIEDLIQSMHIDATLYTESDVHPSCKVTAMKCFLLELQVISLESGDASIHDTVENLIILANNSLSSNGNVTESGCKECEELEEKNIKEFLQSFVHIVQMFINTSAAQELRRIGDEFASPAPAPHPVGLLARVPLSLYSGHPVGLLARVPLSLYSGLSGRSDNHLSGRSDNHPAPAP[Bcl-2] | Rigid linker-(L3)$_3$-rigid linker |
